# Variation in root system architecture among the founder parents of two 8-way MAGIC wheat populations for selection in breeding

**DOI:** 10.1101/2021.07.12.451891

**Authors:** Shree R. Pariyar, Kerstin A. Nagel, Jonas Lentz, Anna Galinski, Jens Wilhelm, Alexander Putz, Sascha Adels, Kathrin Heinz, Claus Frohberg, Michelle Watt

## Abstract

Root system architecture (RSA) is a target for breeding because of the interest to develop crops with roots that use nutrients and water more effectively. Breeding for RSA requires phenotypic diversity in populations amenable to QTL identification to provide markers for large breeding programs. This study examined the variation for root traits across the parents of two multi-parent advanced generation inter-cross (MAGIC) wheat populations from NIAB and CSIRO for 16 days in an upgraded version of the non-invasive, germination paper-based phenotyping platform, *GrowScreen-PaGe*. Across all parents, total root length varied up to 1.90 fold, root biomass 2.25 fold and seminal root angle 1.16 fold. The CSIRO parents grew faster, exhibited slightly wider seminal root angle and produced larger root systems compared to NIAB parents. Lateral root lengths, leaf lengths and biomass contrasted most between fastest (Robigus - NIAB and AC Barrie - CSIRO) and slowest growing parents (Rialto - NIAB and G204 Xiaoyan54 - CSIRO). Lengths of lateral and total root, and leaf number and length had moderate to high heritability (0.30-0.67) and repeatability. Lengths of lateral roots and leaves are good targets for enhancing wheat crop establishment, a critical stage for crop productivity.

## INTRODUCTION

Bread wheat (*Triticum aestivum* L.) is a staple food crop for the world, providing 19% of the calories and 20% of the protein in human diets (FAO, 2017; Shewry, 2009). Wheat is an allohexaploid with three closely-related but independently maintained genomes combined by multiple hybridizations among three progenitor species. The first hybridization occurred between the wild diploid wheat T. urartu (AA, 2n = 14) and an unknown species containing the B genome (BB, 2n = 14, most probably *Aegilops speltoides*), resulting in the tetraploid ancestor of modern wheat species, wild emmer wheat *T. turgidum* ssp. *dicoccoides*, AABB, 2n = 28). Wild emmer further hybridized with goat grass A. tauschii (DD, 2n = 14) to produce today’s modern bread wheat (Dubcovsky and Dvorak, 2007; Huang et al., 2002; Matsuoka, 2011; Shewry, 2009). As a consequence of hybridization of three genomes, the entire bread wheat genome is exceptionally large compared to the other staple cereals maize and rice, and only recently a fully annotated reference genome has become available (Appels et al., 2018).

Genetic improvement of wheat relies on diversity in the phenotypes and genotypes of parents and populations, and heritable and repeatable quantitative trait loci (QTL). We analyzed the shoot and root phenotypic variation of two sets of eight way MAGIC wheat parents: CSIRO MAGIC (Huang et al., 2012) and NIAB MAGIC (Mackay et al., 2014) (Figure 1). MAGIC mapping populations have high allelic diversity, high levels of recombination, and the genomes form a fine-scale mosaic of all the parents (Cavanagh et al., 2008; Huang et al., 2011; Kover et al., 2009). Combined with high density molecular markers and sequence information, these populations serve as a new generation of mapping populations for QTL discovery. Several populations are available for wheat (Cavanagh et al., 2008; Fradgley et al., 2018; Huang et al., 2012; Huang et al., 2015; Mackay et al., 2014; Stadlmeier et al., 2018). The eight CSIRO MAGIC parents are spring wheats from Australia, Asia, and North America, selected based on diverse geographical distribution and agronomical performance (Huang et al., 2015). These parents were previously used to identify QTL for hair length (Delhaize et al., 2015), above ground coleoptile and seedling growth (Rebetzke et al., 2014), spikelet formation and grain dormancy (Barrero et al., 2015; Boden et al., 2015) and canopy architecture (Richards et al., 2019). The NIAB MAGIC parents include eight european winter wheat from the United Kingdom and France (Figure 1), and were selected based on yield, bread-making quality and disease resistance (Mackay et al., 2014). The NIAB MAGIC wheat populations were successfully used to identify QTLs linked to leaf senescence and plant height (Camargo et al., 2016), presence or absence of awns (Mackay et al., 2014), resistant to *Septoria tritici* (Riaz et al., 2020), and leaf and glume blotch (Lin et al., 2020). To date only one study reports the use of a wheat MAGIC population to identify genetic variation in root traits. Delhaize et al. (2015) identified major QTLs linked to rhizosheath size (using bound soil and root hair length as phenotypes) in the CSIRO MAGIC wheat population.

**Figure 1.**
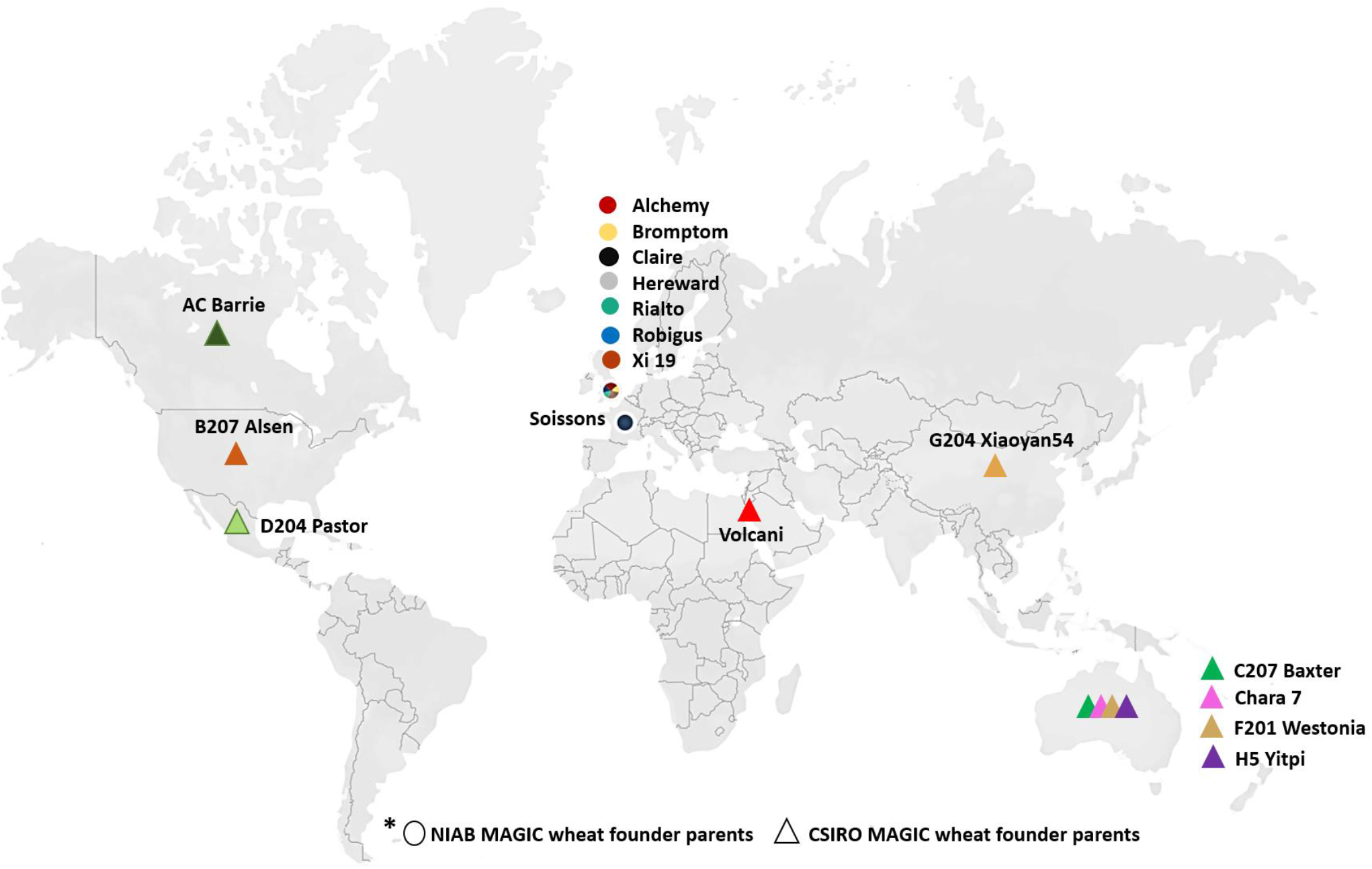
Geographic distribution of MAGIC wheat founder parents used in this study. NIAB MAGIC founder parents (circle symbols) originated from the UK (7) and France (1). The CSIRO MAGIC founder parents (triangle symbols) originated from Australia (4 lines) and one each from China, Israel, Canada, USA, and Mexico. The details of the origin and attributes of both set of MAGIC founder parent lines are listed in Table S1.

Improving root traits for breeding is a promising strategy to speed up genetic gain for plant performance (Tracy et al., 2019). Root system architecture (RSA) is the spatial distribution of roots in a particular environment and is determined by genetic and environmental factors (Lynch, 1995; Rich and Watt, 2013). Underlying components of RSA are root number, elongation rate and root angle (Rich and Watt, 2013); these measures can result in root lengths with depth through the soil (Asif and Kamran, 2011; Carvalho et al., 2014; King et al., 2003), including in the early vegetative phase in different field conditions (Rich et al., 2020). In wheat and other small grain cereals, root systems are composed of root types, differentiated based on their tissue of origin. A seminal root system originates from the seed embryo, and a nodal root system originates from leaf nodes, each with axile roots and lateral roots. The seminal roots are the first to emerge from the seed upon germination, and their elongation rate and angle support axial transport of water and nutrients to the shoot until at least 3 to 4 leaves, when nodal axile roots emerge if conditions are favorable (Watt et al., 2008). Lateral roots begin to make up most of the root length of the entire root system after two leaves across a range of species including wheat (Gioia et al., 2016). In maize they have been shown to be important for accessing water (Ahmed et al., 2016; Tracy et al., 2019), at least in part because of their dominant length (Varney and Canny, 1993). It is important to use a phenotyping method for roots that allows quantification of all components of RSA, including the fine lateral roots.

Phenotypes for QTL must be repeatable and heritable (Richards et al., 2010; Wissuwa et al., 2016). Controlled environment phenotyping is currently a preferred option to field phenotyping because of the ability to control variability and increase repeatability and heritability (Watt et al., 2020). The phenotypes of young wheat plants such as seminal root length and total root length measured in control environments correlate well to expression in fields of those phenotypes (Watt et al., 2013, 2020; Rich et al., 2020). Direct root phenotyping in the field is difficult mainly due to the opaque nature of soil, and is restricted to coring, shovelomics and minirhizotrons, which only reveal part of root systems (Tracy et al., 2019; Wasson et al., 2020). Here, we used the non-invasive, high-throughput phenotyping system ‘*GrowScreen-PaGe*’, previously applied to shoots and roots of young rapeseed, maize, barley, and wheat plants (Gioia et al., 2016). This system has several features critical to root phenotyping: precision, as it allows the complete root system to be quantified in 2D using a high resolution camera; repeatability, because plants are grown on germination paper to control for effects of soil variability; and throughput, as the method can scale to high replicates and plant numbers, and be readily established in other laboratories.

This study reports on the phenotypic variation in early root and shoot traits of the CSIRO and NIAB MAGIC wheat parents. By using *GrowScreen-PaGe*, we quantify the variation, heritability and repeatability of component phenotypes of RSA including seminal and lateral lengths, angles and dynamic traits such as rates of growth, extracted from images. We also determined allocation between root and shoot biomass within and across the MAGIC wheat parents. The aim of the study was to identify root traits in these parents of MAGIC populations that could be selected in future pre-breeding and breeding using phenotyping or QTL-based markers.

## MATERIALS AND METHODS

### Plant Materials

The founder parents for the NIAB and CSIRO wheat MAGIC populations have diverse origins (Figure 1), pedigrees, yield, grain quality, disease resistance and growth parameters (Table S1). The NIAB parents (Mackay et al., 2014) are winter wheat cultivars from the United Kingdom (Alchemy, Brompton, Claire, Hereward, Rialto, Robigus and Xi 19) and France (Soissons) (Figure 1). The CSIRO parents (Huang et al., 2012) are spring wheat cultivars from Australia (Yitpi, Baxter, Chara7 and Westonia) and China (G204 Xiaoyan54), Israel (Volcani), Canada (AC Barrie), USA (B207 Alsen), and Mexico (D204 Pastor) (Figure 1). The cultivar Chara7 was included to develop 4-way population, however was excluded from 8 way MAGIC wheat population (Table S1).

### Phenotyping System

Phenotyping was performed using the *GrowScreen-PaGe* phenotyping platform which is based on repeated imaging of root development on wetted germination paper. The platform used is as described in Gioia et al. (2016) with the following modifications: wetted germination paper (smooth dark blue 194 grade paper; Ahlstrom Germany GmbH, Bärenstein, Germany) was fixed onto dark grey PVC plates (RAL 7011, Max Wirth GmbH, Germany) rather than transparent PMMA plates with transparent shirt clips (Hemdklammer 38mm, Georg Scharf GmbH, Balingen, Germany); new opaque polypropylene containers (60 × 40 × 42 cm, Eurobehälter, EG 64/42 HG, Auer Packaging GmbH, Germany) replaced those used in Gioia et al. (2016) to prevent light entering through handles; PVC containers were covered by a new custom-made polypropylene lid with adjustable black strip brushes in the middle that allow leaf emergence and growth (Mink GmbH & Co. KG, Germany); and PVC plates (37.5 × 30 × 0.2 cm) and germination paper (37 × 25 cm) were adjusted in size to fit the new containers (see Supplementary Figure S1A, B for the new set up).

### Plant Cultivation and Experimental Design

Seeds were pre-coated with the fungicide prothioconazole (33 ml/100 kg seed) and then germinated in the dark in Petri dishes between two wet filter papers at 22 °C for 24 h. Seeds with the first seminal root emerged from the base of the embryo and ca. 1.5-1.7 mm length were transplanted onto the *GrowScreen-PaGe* germination paper with the seminal root facing downward. The seedlings were fixed to the germination paper using a transparent, self-adhesive tape (Opsite Flexifix, 7478029, Smith & Nephew GmbH) (Figure 2), replacing the fold-back butterfly clips and filter paper used in Gioia *et al*. (2016), allowing the angle of seminal root emergence to be imaged. After fixing the germinated seed, a modified Hoagland nutrient solution (Hoagland and Arnon, 1950) was sprayed evenly over the germination paper to the point of saturation with Gloria sprayers (Gloria Haus & Gartengeräte GmbH, Germany, EAN 400 632 571 7872). Up to 25 PVC plates with one germination paper and one seedling on each side (two plants per plate) were positioned vertically and 2.0 cm apart in the opaque containers (Supplementary Figure S1B). Each container was filled with 12 L of the modified Hoagland nutrient solution, submerging the bottom 5 cm of germination papers. The nutrient solution was replaced once a week. Each experiment contained 8 replicates (plants) of each parent (8 plants x 17 parents = 136 plants) which were arranged in a completely randomized block design. In total, three successive experimental repeats were conducted. Plants were grown in a controlled growth chamber at 22°C/18°C (day/night) air temperature, 12 h/12 h light/dark, 60% relative air humidity and ~150 μmol m^−2^ s^−1^ photosynthetically active radiation (PAR) at leaf level (Johnson Controls Systems & Service GmbH, Leipzig, Germany).

**Figure 2.**
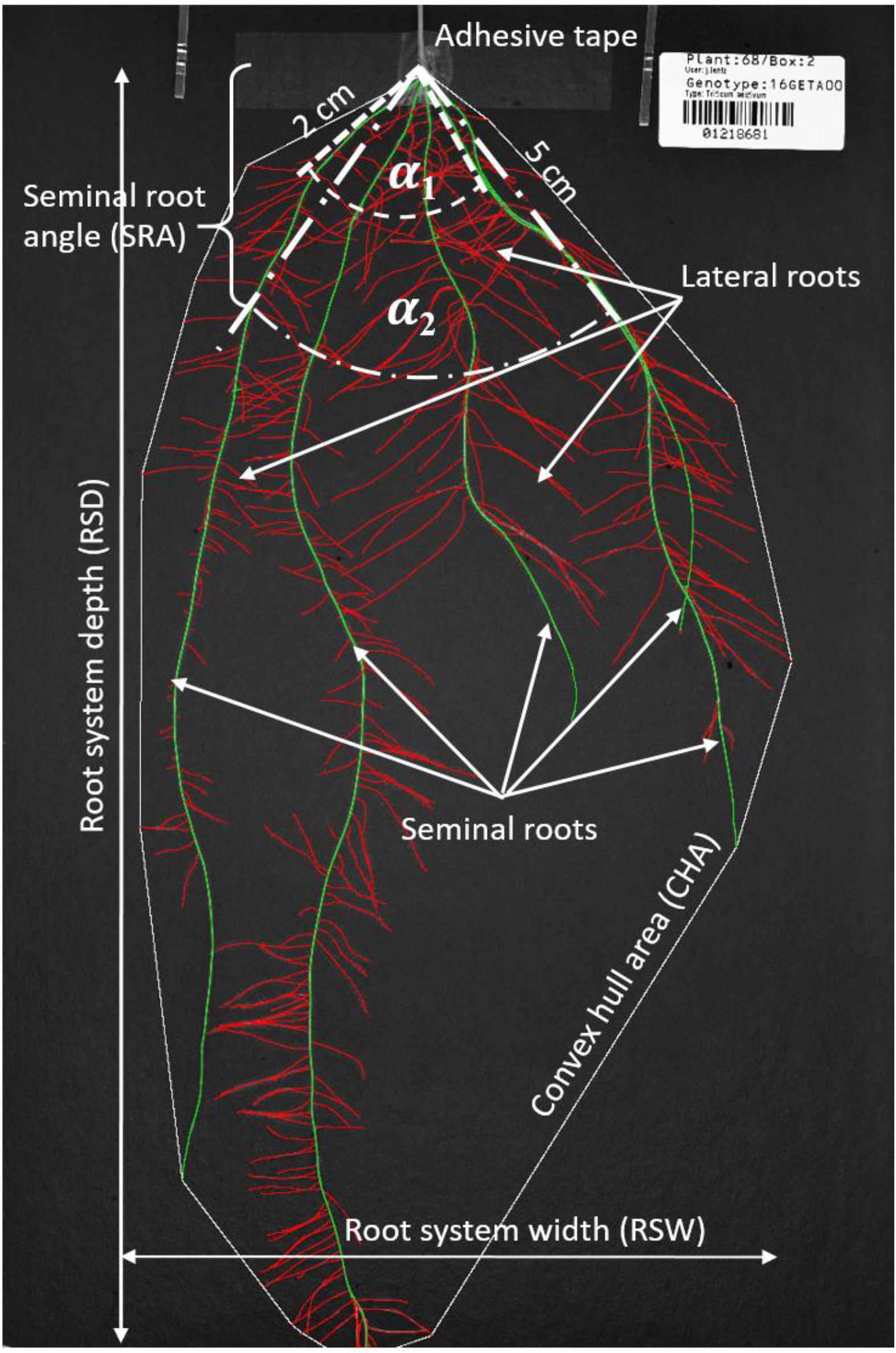
Typical wheat root system analyzed at 16 days after transplanting by the image processing workflow of the *GrowScreen-PaGe* platform. Root lengths of different root classes (seminal roots marked in green; and lateral roots marked in red) are analyzed separately. Furthermore spatial information, such as max. root system depth (RSD) and width (RSW) (indicated by white vertical and horizontal lines) as well as convex hull area (CHA), the area covered by the root system indicated by the white line surrounding the root system are quantified. The seminal roots angle (SRA) between the two outermost seminal roots was measured at 2 (*α*_1_) and 5 cm (*α*_2_) distances from the seed.

### Image Acquisition and Analysis

Root systems were photographed at 2, 9 and 16 days after transplanting (DAT) onto the germination papers, using the mobile imaging box described in Gioia *et al*. (2016) with an upgraded, less manual image-processing workflow. Three major software parts and steps were added. First is the ‘Segmenter’, which segments root parts from the background and labels those as developmental types based on their points of origin (seminal roots originating from seed; lateral roots originating from seminal or first order lateral roots). The second step is called ‘PaintRoot’, a graphical user interface to check the automatic labeling provided by the ‘Segmenter’ and allows manual correction of the segmentation and labeling in the mask images if required. The third step is the ‘Analyzer’, which processes the checked or corrected image (Supplementary Figure S1D) to quantify the root phenotypes. These new software steps were used to quantify total root length (TRL), seminal root length (SRL), lateral root length (LRL), seminal root angle (SRA) between two outermost seminal roots at a distance of 2 cm (SRA_2cm_) and 5 cm from the seed (SRA_5cm_), root system depth (RSD), root system width (RSW), and convex hull area (CHA). See detailed list and definition of all measured root and shoot growth traits in Table 1 and Figure 2.

**Table 1.** List of measured root and shoot traits in this study.

| Trait | Description | Unit |
| --- | --- | --- |
| First seminal root (FSR) | Length of radicle (measured 1 day after germination) | mm |
| Seminal root length (SRL) | Length of all seminal roots | cm |
| Lateral root length (LRL) | Length of all roots branched from seminal roots | cm |
| Total root length (TRL) | Total sum of seminal and lateral root length | cm |
| Root system depth (RSD) | Maximal vertical depth of a root system | cm |
| Root system width (RSW) | Maximal horizontal distribution of a root system | cm |
| Convex hull area (CHA) | Area of the convex hull that encompasses the root system | cm<sup>2</sup> |
| Seminal root angle (SRA) | Angle between the outermost left and right seminal roots measured at 2 or 5 cm distance from seed respectively | ° |
| Leaf length (LeafL) | Length between the seed and tip of the longest leaf | cm |
| Number of leaves (LeafNo) | Number of leaves (without cotyledons) | - |
| Root dry weight (RDW) | Root dry weight | g |
| Shoot dry weight (SDW) | Shoot dry weight | g |
| Root mass ratio (RMR) | Dry mass of root divided by the total dry mass of entire plant | g/g |
| Shoot mass ratio (SMR) | Dry mass of shoot divided by the total dry mass of entire plant | g/g |

Plants were harvested at 16 DAT and number of leaves and length of longest leaf were measured manually. Further, roots and shoots were separated above the seed, oven dried at 65°C for one week and dry weight was recorded using Mettler Toledo™ (Model: XS-205DU, analytical balances, Fischer Scientific).

### Statistical Analysis

The data were analyzed with a one-way analysis of variance (ANOVA), noting significant differences between means at a level of *P ≤ 0.05* using SigmaPlot 13.0 (Sigma Stat, Systat Software Inc., Richmond, CA, USA). The coefficient of variation (CV) was calculated as the ((SD/mean)*100), where SD = standard deviation. The CV indicates the extent of variation of a phenotype relative to its mean. Broad-sense heritability (*H^2^*) was calculated in SAS (SAS 9.4) according to (Falconer and Mackay, 1996; Holland *et al*., 2003):

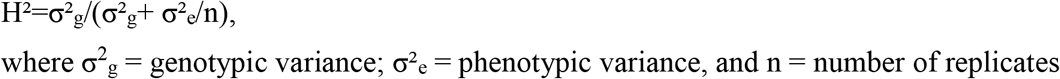

Repeatability (*R*) across three successive experiments was calculated as the variance among group means (group-level variance *V_G_*) over the sum of group-level and data-level (residual) variance *V_R_* followed by confidence interval at 1000 bootstrap (Stoffel et al., 2017). Pearson correlations and regressions between traits were evaluated by generating a correlation matrix. Repeatability, correlations and regression were analyzed in R3.5 (RCore, 2016).

## RESULTS

### Variation in Root Lengths and Biomass was Observed in Both sets of MAGIC Parents

The parents of both MAGIC wheat populations had substantial variability for root length and dry weight phenotypes at the final time point (16 DAT) of the study (Figure 3). Within the NIAB parents, Robigus, Bromptom and Alchemy had the longest median total root length while Rialto and Hereward cultivars had the shortest median total root length (Figure 3A). Within the CSIRO parents, AC Barrie, B207 Alsen, and D204 Pastor had the longest median total root length while G204 Xiaoyan54 and C207 Baxter had the shortest (Figure 3A). Root dry biomass ranged from 17 to 27 mg and 18 to 33 mg for the NIAB and CSIRO parents, respectively (Figure 3D). On average, the CSIRO parents had 1.3 times greater total root length than the NIAB parents. This was due to slightly longer (1.1 times) seminal roots and relatively longer (1.3 times) lateral roots. CSIRO founder parents also had greater root as well as shoot biomass (1.2 times) than the NIAB parents (Figure 3D, Supplementary Tables S1 and S2). Additionally, we found 41% longer leaf length and 40% more shoot dry weight with CSIRO parents compared to NIAB (Supplementary Tables S2 and S3).

**Figure 3.**
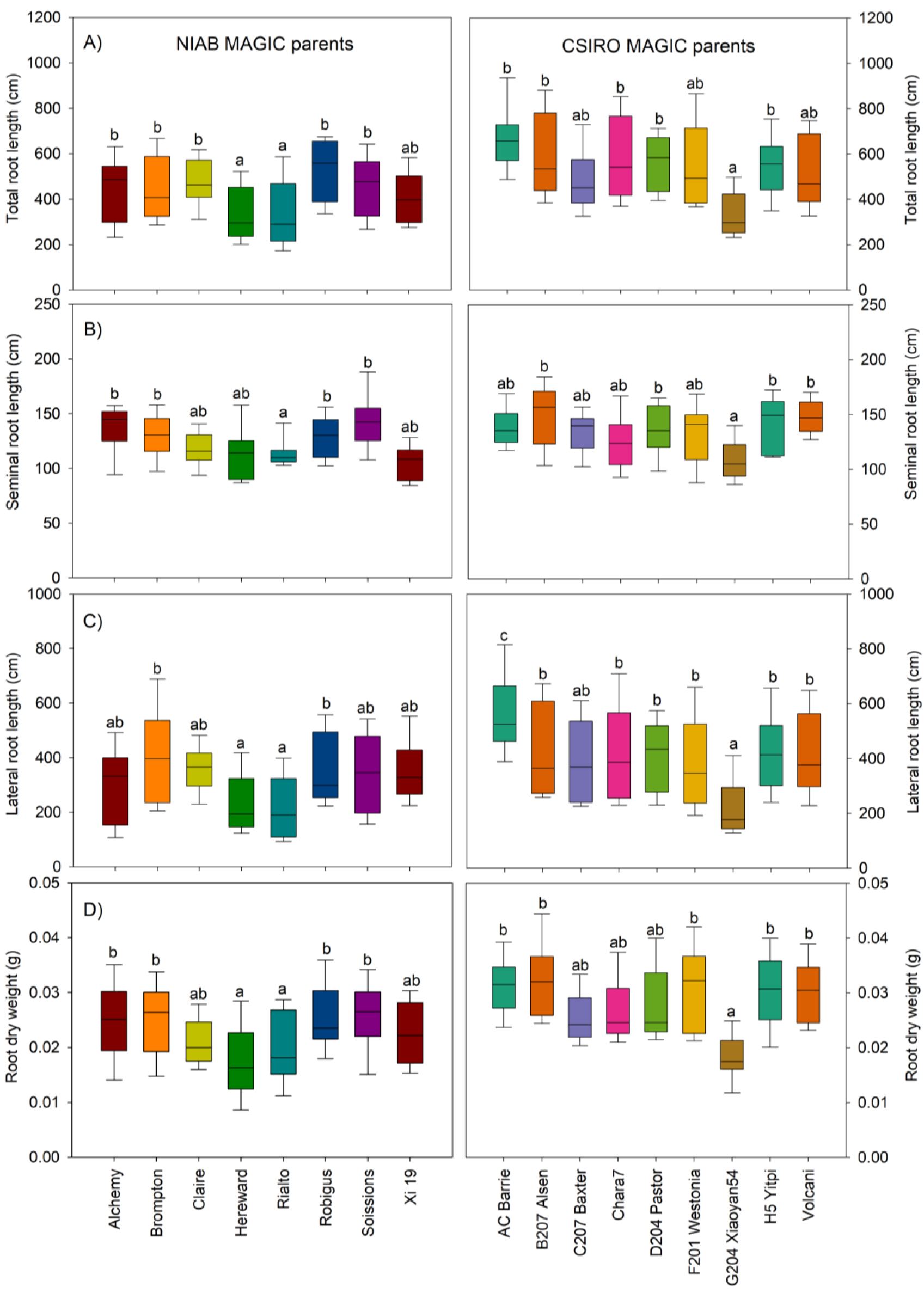
Phenotyping variation in root traits measured at 16 days after transplanting (DAT). NIAB MAGIC wheat founder lines (left panel) and CSIRO MAGIC wheat founder parents (right panel). (A) Total root length, B) Seminal root length, C) Lateral root length, and D) root dry weight. The boxes indicate the 25^th^ and 75^th^ percentile of the distribution, the ‘whiskers’ the 10^th^ and 90^th^ percentile and the lines in the middle of the box the median value. Boxes with different letters indicate significant differences at *P < 0.05* according to one way ANOVA analysis, a≤ b ≤ c, n=24 (pooled data from 3 successive experiments).

### Slow and Fast growing Founder Parents Differed most through Elongation rates of Lateral Roots

We dissected the temporal dynamics and root types for the two parents per population that contrasted most for total root length (Figure 4). By 16 DAT, within NIAB founder parents, the slowest growing Rialto had 1.5 fold less total root length than the fastest growing Robigus (Figure 4A, B and C); within CSIRO parents, the slowest growing G204 Xiaoyan54 had 1.9 fold less total root length than AC Barrie (Figure 4E, F and G). Lateral root length variation was most notable at driving these contrasts, differing 1.7 times between the slowest and fastest parents of the NIAB parents, and 2.2 times between the two extreme CSIRO parents at 16 DAT. Further, analysis of root growth dynamics over time showed that rates of lateral root growth were higher in fastest growing parents compared to slow growing parents in both NIAB and CSIRO MAGIC parents (Figure 4D, H). The growth rate of lateral roots was 2.6 times higher in Robigus compared to Rialto, while lateral root growth rate in AC Barrie was 3.4 times higher compared to G204 Xiaoyan54. However, no significant differences were recorded for root system depth between fastest and slowest growing parents although the fast growing parents grew slightly deeper (Figure 4I, J). At 16 DAT, most of the roots had reached the bottom of the germination paper, restricting extraction of reliable data at this imaging time point.

**Figure 4.**
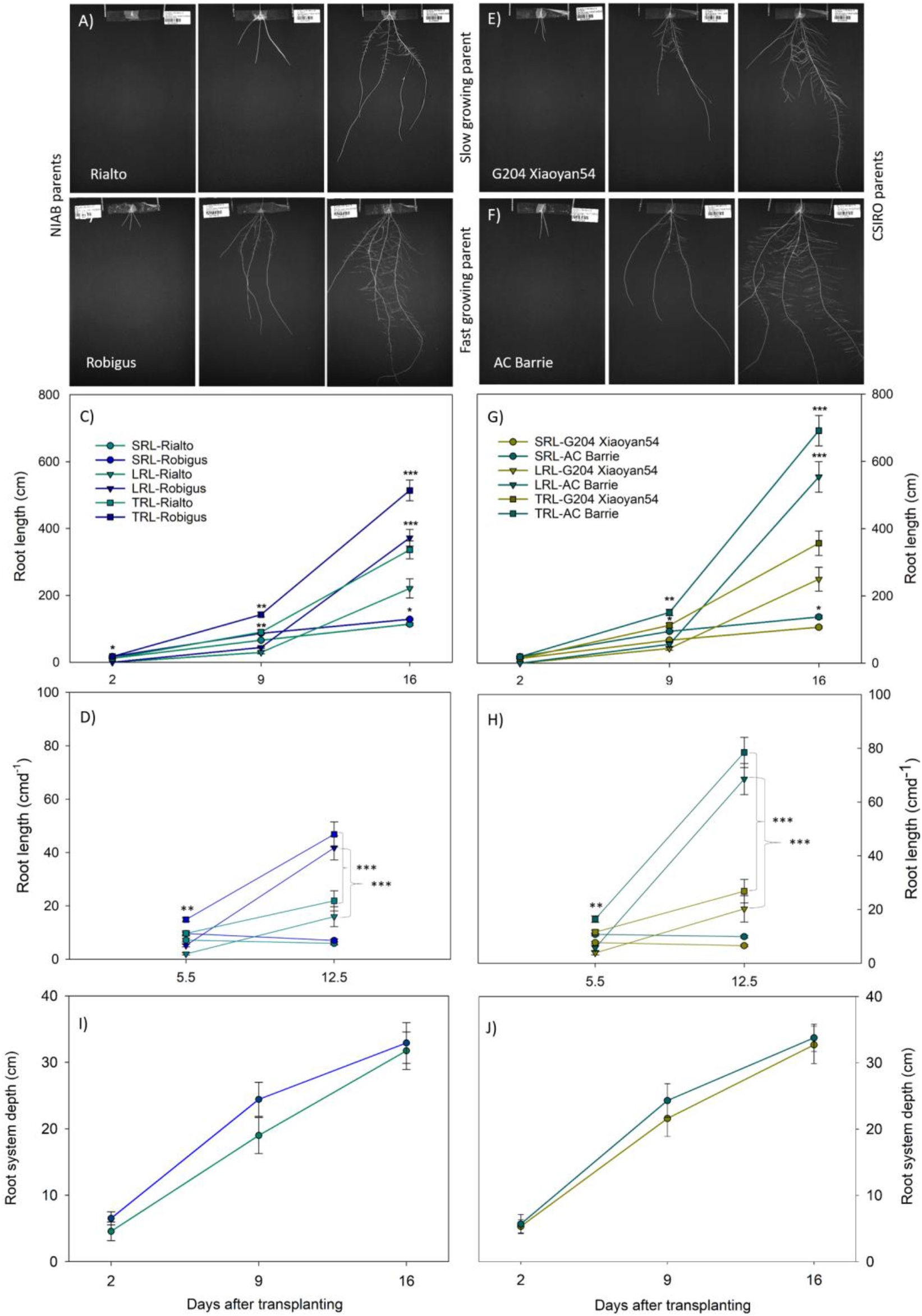
Root growth dynamic of selected slow and fast growing MAGIC wheat founder parents over time. The slow growing lines Rialto (NIAB) and G204 Xiaoyan54 (CSIRO) and fast growing lines Robigus (NIAB) and AC Barrie (CSIRO) were selected based on total root length differences at 16 DAT. Typical original root images of slow (A, D) and fast growing lines (B, E) at 2, 9 and 16 days after transplanting. Seminal root length (SRL), lateral root length (LRL), and total root length (TRL) (C, G), dynamic of root growth (D, H) and root system depth (I, J) were analyzed. Line with asterisk indicate significant differences at (\**p < 0.05*, \*\**p < 0.01*, \*\*\**p < 0.001*, n=24) according to one way ANOVA analysis.

### Phenotypic Variation for Seminal Root Angles was Observed only Across CSIRO MAGIC Parents

We analyzed the root angle between the two widest seminal roots at SRA_2cm_ and SRA_5cm_, and found significant variation only across CSIRO MAGIC parents (Figures 5, S2), although SRA_2cm_ and SRA_5cm_ increased and were correlated positively through time (see Supplementary Figure S2, Tables S2 and S3). No significant differences for seminal root angle were found within NIAB parents (Figure 5A, B). NIAB parents had a slightly narrower SRA_2cm_ compared to that of CSIRO over time (Figure S2 A-C), but SRA_5cm_ was narrower in CSIRO parents than in NIAB (Figure 5A, B). For CSIRO MAGIC parents, wide SRA_5cm_ correlated to the total length and rate of total root length growth of the root system (Figures 5A, B, C). However, this pattern of wider root angle linked to fast growing parent (AC Barrie) and narrow angle to slow growing parent (G204 Xiaoyan54) was not found for NIAB parents (Figures 5A, B, S2 A-C).

**Figure 5.**
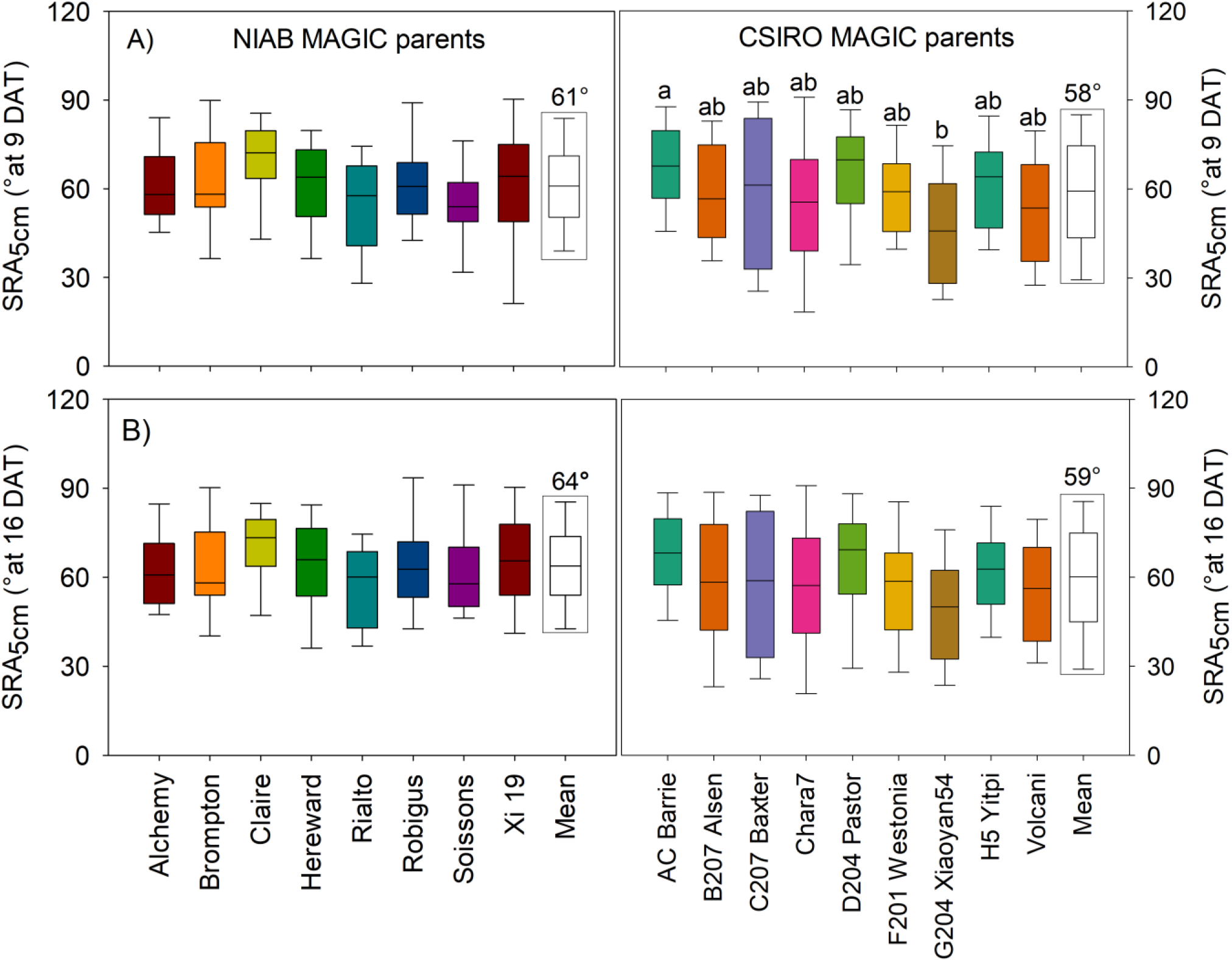
Variation for root angle measured between the two outmost seminal roots at a distance of 5 cm (SRA_5cm_) over time. A) SRA_5cm_ at 9 days after transplanting (DAT), and B) SRA_2cm_ at 16 DAT. The boxes indicate the 25^th^ and 75^th^ percentile of the distribution, the ‘whiskers’ the 10^th^ and 90^th^ percentile and the lines in the middle of the box the median value. No significant differences were recorded at *P < 0.05*, n=24 according to one way ANOVA analysis. Boxes highlighted with rectangular black line indicates mean seminal root angle (SRA) across NIAB and CSIRO wheat founder parents.

Our results suggested that variation in seminal root angles (SRA_2cm_ and SRA_5cm_) depended on cultivar, and time and position of the measurement (Figure 5, Supplementary Figure S2). Note that at 2 DAT, the length of seminal roots were shorter than 5 cm and consequently no SRA_5cm_ could be quantified, and was measured at 9 and 16 DAT only (Supplementary Tables S2 and S3).

### Traits with High Heritability, Repeatability and Correlations for Selection in Breeding

Heritability (H^2^) and repeatability (*R^2^*) ranged widely for root and shoot traits in both sets of MAGIC wheat parents (Table 2). Heritability was highest at 0.67) for NIAB, and 0.84) for CSIRO parents (Table 2). Both sets of parents had high heritability for leaf length and number (Table 2); moderate heritability for first seminal, lateral and total root length; and low heritability for root system depth, root system width, seminal root angle at 2 and 5 cm distance, and convex hull area. Repeatability was highest for total root length (0.67) and lowest (0.10) for root system depth across MAGIC parents (Table 2). Repeatability was also low for root system width, seminal root angle and convex hull area; while moderate to high repeatabilities were recorded for seminal and lateral root length, and leaf length (Table 2).

**Table 2.** Heritability and repeatability of root and shoot traits in both sets of MAGIC wheat parent parents. Heritability was analyzed between NIAB and CSIRO parental parents separately while repeatability was analyzed by combining both NIAB and CSIRO parents (n=24). *DAG day after germination, DAT day after transplanting, low (≤ 0.3), moderate (0.3 - 0.6) and high (> 0.6).

| Traits | Heritability ($h^2$) | | Repeatability ($R^2$) |
| --- | --- | --- | --- |
|  | MAGIC founder |  | MAGIC founder parents |
|  | NIAB | CSIRO |  |
| First seminal root (1 DAG) | 0.43 | 0.64 | 0.36 |
| Seminal root length (2 DAT) | 0.29 | 0.32 | 0.58 |
| Seminal root length (9 DAT) | 0.20 | 0.44 | 0.54 |
| Seminal root length (16 DAT) | 0.23 | 0.35 | 0.44 |
| Lateral root length (9 DAT) | 0.40 | 0.22 | 0.34 |
| Lateral root length (16 DAT) | 0.36 | 0.38 | 0.65 |
| Total root length (2 DAT) | 0.29 | 0.32 | 0.58 |
| Total root length (9 DAT) | 0.36 | 0.19 | 0.44 |
| Total root length (16 DAT) | 0.35 | 0.40 | 0.67 |
| Root system depth (2 DAT) | 0.38 | 0.07 | 0.30 |
| Root system depth (9 DAT) | 0.30 | 0.19 | 0.42 |
| Root system depth (16 DAT) | 0.12 | - | 0.10 |
| Root system width (2 DAT) | 0.25 | 0.12 | 0.24 |
| Root system width (9 DAT) | 0.24 | 0.30 | 0.34 |
| Root system width (16 DAT) | 0.16 | 0.29 | 0.23 |
| Seminal root angle2cm (2 DAT) | - | 0.06 | 0.23 |
| Seminal root angle2cm (9 DAT) | 0.06 | 0.10 | 0.14 |
| Seminal root angle2cm (16 DAT) | - | 0.22 | 0.06 |
| Seminal root angle5cm (9 DAT) | 0.41 | 0.10 | 0.08 |
| Seminal root angle5cm (16 DAT) | 0.30 | 0.20 | 0.21 |
| Convex hull area (2 DAT) | 0.39 | 0.13 | 0.33 |
| Convex hull area (9 DAT) | 0.23 | 0.24 | 0.32 |
| Convex hull area (16 DAT) | 0.12 | 0.26 | 0.19 |
| Leaf length (16 DAT) | 0.60 | 0.84 | 0.58 |
| Leaf number (16 DAT) | 0.67 | 0.61 | 0.42 |
| Root dry weight (16 DAT) | 0.25 | 0.50 | 0.34 |
| Shoot dry weight (16 DAT) | 0.37 | 0.50 | 0.38 |

Similarly, correlations between root types and shoot phenotypes ranged widely in both sets of MAGIC parents (Figure 6A, B). At 16 DAT, high correlation were recorded between root dry weight and shoot dry weight (*R^2^*=0.775 for NIAB and *R^2^*=0.741 for CSIRO, Figure 7D) while the lowest correlations (*R^2^*=0.002-0.007 Figure 6A, B) were recorded between seminal root angle and root system depth, and seminal root angle and root system width in both MAGIC sets. For NIAB parents, total root length and root dry weight (*R^2^*=0.528, Figure 7B), and total root length and shoot dry weight (*R*^2^=0.516, Figure 7C) were moderately correlated, and seminal and lateral root length (*R^2^*=0.22 for NIAB and *R^2^*=0.10 for CSIRO, Figure 7A) had low correlations. For CSIRO parents, correlations were slightly higher than those of NAIB (*R^2^*=0.692 - total root length vs. root dry weight, Figure 7B and *R^2^*=0.620 - total root length vs. shoot dry weight, Figure 7C).

**Figure 6.**
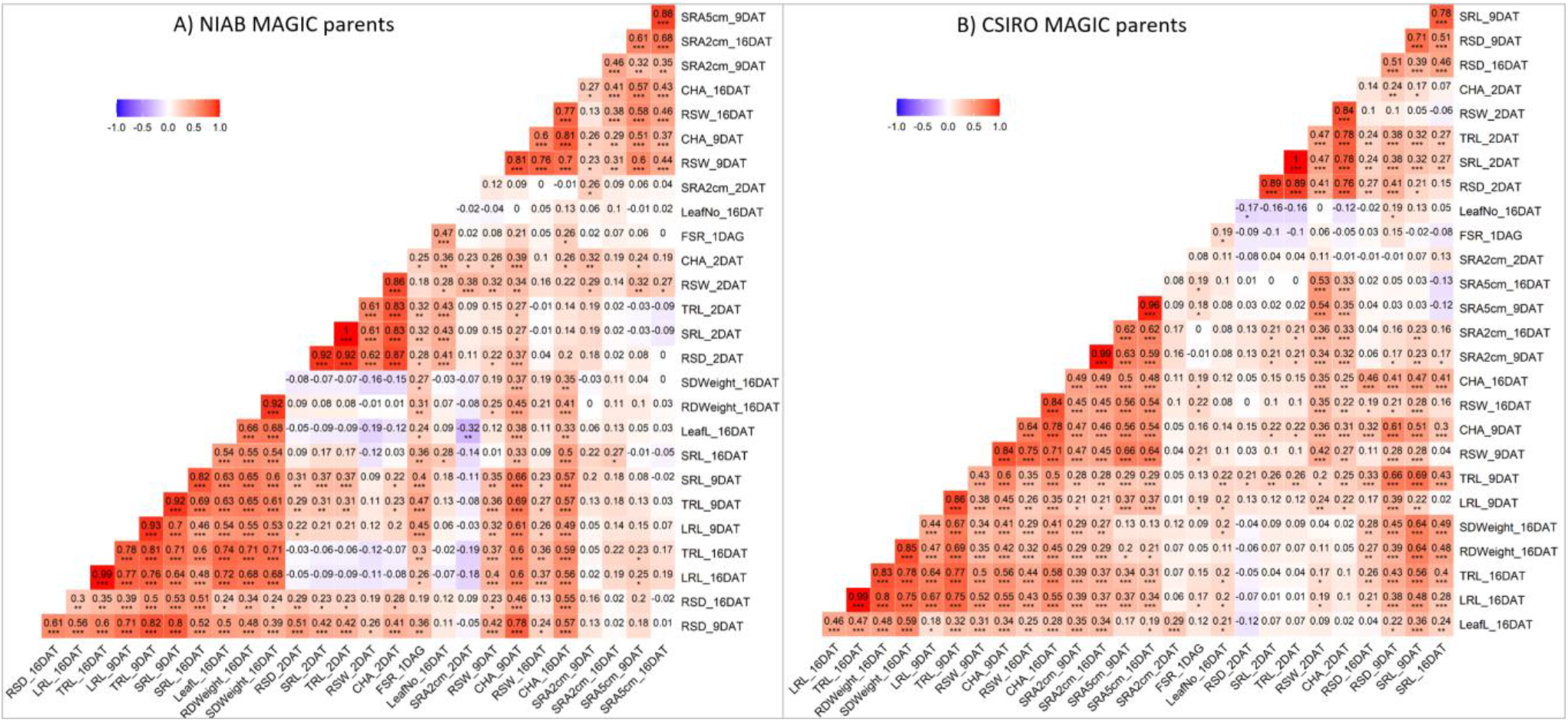
Pearson’s correlation matrix between root and shoot traits of MAGIC wheat founder parents, A) NIAB MAGIC wheat parents and B) CSIRO MAGIC wheat parents. Intensity of color indicates correlation among the traits ranges from (1 to −1) and is the fraction of the variance in the two variables that is “shared”. FSR first seminal root length, SRL seminal root length, LRL lateral root length, TRL total root length, SRA seminal root angle, RSD root system depth, RSW root system width, CHA convex hull area, RDWeight root dry weight, LeafL longest leaf length, LeafNo leaf number, SDWeight shoot dry weight, DAG days after germination, DAT days after transplanting. Number with asterisk indicates significant differences at \**p < 0.05*, \*\**p < 0.01*, \*\*\**p < 0.001* and *n*=24.

**Figure 7.**
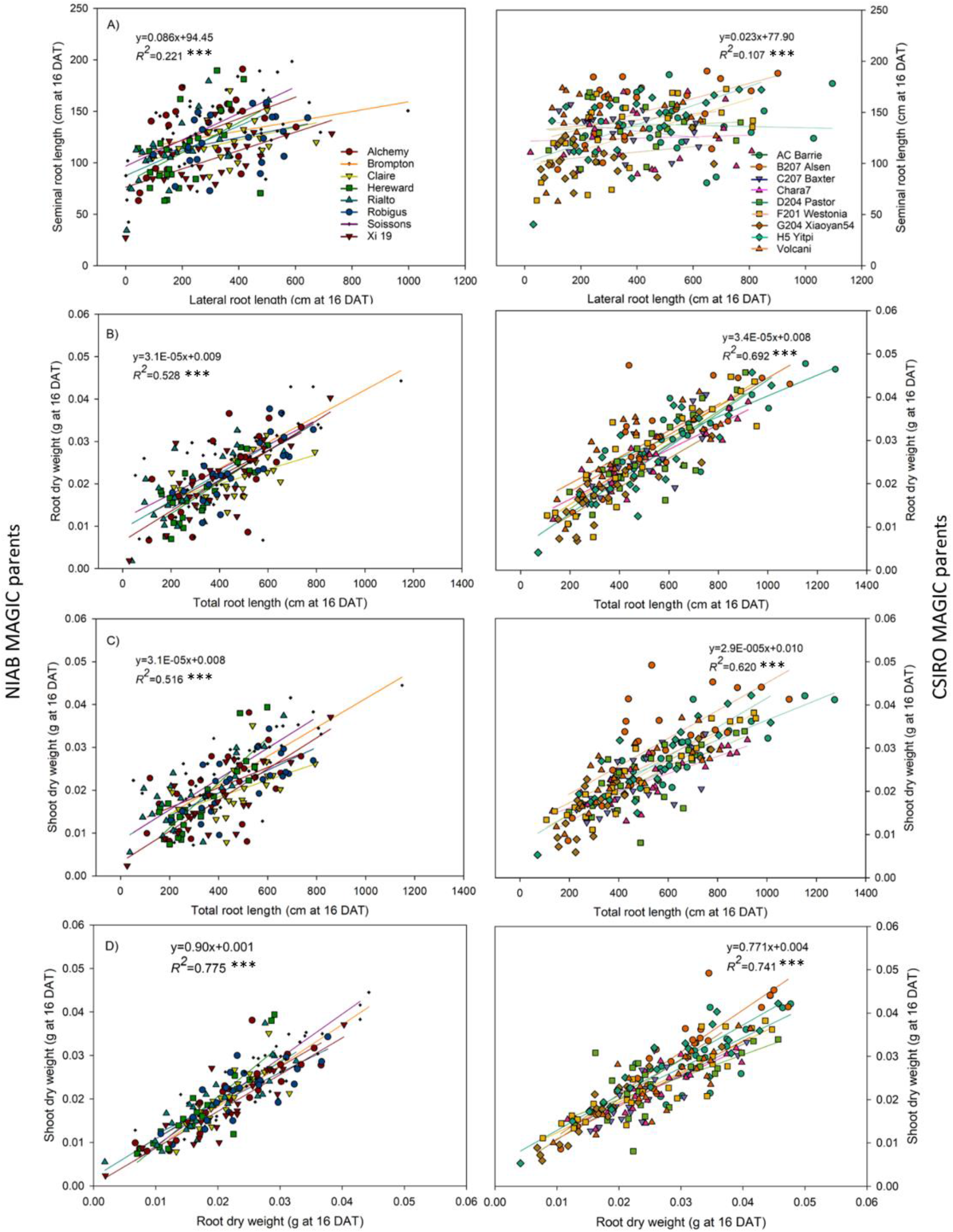
Relationship between traits of MAGIC wheat founder parents (left panel - NIAB, and right panel - CSIRO) at 16 days after transplanting (DAT). A) correlation between seminal and lateral root length, B) correlation between root dry weight and total root length, C) correlation between shoot dry weight and total root length, and D) correlation between shoot and root dry weight. Mean of correlations across MAGIC founder parents were shown (n=24, \*\*\**p < 0.001*).

Although, heritability, repeatability and correlations between root and shoot traits varied over time, lengths of first seminal root, lateral root and total root lengths, and leaf number and length had the most consistent moderate to high heritability and repeatability. These traits can be targeted for selection in breeding. CV for all root and shoot phenotypes was as high as 62% across NIAB and as high as 63% for CSIRO parents (Supplementary Table S5 and S6). Lower CVs (< 40%) were recorded for root system depth and width, while higher CVs (≥40%) were linked to first seminal root length, lateral root length and convex hull area of both sets of MAGIC parents.

### Fast Growing Parents Allocated More Resources to the Roots during Early Stages of Plant Development

Root and shoot biomass allocation varied between and across both sets of MAGIC parents at 16 DAT (Table 3). CSIRO parents had 1.2 fold greater root and shoot dry biomass than NIAB parents (Table 3). Fast growing parents produced more total plant biomass (Robigus produced 1.22 fold more biomass compared to Rialto, while AC Barrie produced 1.7 fold more compared to G204 Xiaoyan54) compared to slow growing parents in both sets of MAGIC parents (Tables S2, S3). We found striking contrast for root mass ratio (RMR) and shoot mass ratio (SMR) between fast and slow growing parents in both MAGIC sets at 16 DAT. For example, RMR was higher in fast growing parents Robigus (0.52) and AC Barrie (0.53) at 16 DAT while SMR is higher for slow growing parents Rialto (0.51) and G204 Xiaoyan54 (0.50) (Table 3). This reflects that the higher proportion of resources were allocated to the roots during the early stages of plant development in fast growing parents while slow growing parents tend to allocate more resources into shoot development.

**Table 3.** Biomass allocation between root and shoot of both sets of MAGIC wheat parent parents at 16 days after transplanting, mean values are shown. Different letters indicate significant differences (*p=0.05*, n=24; a < b).

| MAGIC wheat | Founder parents | Root dry weight (g) | Shoot dry weight (g) | Root mass ratio | Shoot mass ratio |
| --- | --- | --- | --- | --- | --- |
| NIAB | Alchemy | 0.022b | 0.021ab | 0.513 | 0.487 |
|  | Brompton | 0.027b | 0.024b | 0.525 | 0.475 |
|  | Claire | 0.021ab | 0.020ab | 0.516 | 0.484 |
|  | Hereward | 0.017a | 0.018a | 0.493 | 0.507 |
|  | Rialto | 0.019a | 0.019a | 0.492 | 0.508 |
|  | Robigus | 0.024b | 0.022ab | 0.516 | 0.484 |
|  | Soissons | 0.024b | 0.023b | 0.511 | 0.489 |
|  | Xi 19 | 0.021ab | 0.018a | 0.529 | 0.471 |
| CSIRO | AC Barrie | 0.033b | 0.029b | 0.529 | 0.471 |
|  | B207 Alsen | 0.030b | 0.031b | 0.493 | 0.507 |
|  | C207 Baxter | 0.024ab | 0.021a | 0.53 | 0.47 |
|  | Chara7 | 0.026ab | 0.023ab | 0.533 | 0.467 |
|  | D204 Pastor | 0.027ab | 0.023ab | 0.526 | 0.474 |
|  | F201 Westonia | 0.026ab | 0.024ab | 0.524 | 0.476 |
|  | G204 Xiaoyan54 | 0.018a | 0.018a | 0.497 | 0.503 |
|  | H5 Yitpi | 0.026ab | 0.026ab | 0.501 | 0.499 |
|  | Volcani | 0.028b | 0.024ab | 0.533 | 0.467 |

## DISCUSSION

We aimed to identify root traits in the MAGIC NIAB and CSIRO founder parents that could be selected using phenotyping or QTL-based markers in future. We revealed variation that was heritable and repeatable for component phenotypes contributing to RSA in both sets of parents using the *GrowScreen-PaGe* phenotyping platform. The two traits with greatest heritability and / or repeatability were lateral root length and leaf length (Table 3). We show that measuring RSA non-invasively over time using a high throughput phenotyping platform enables the selection of key functional traits that could be amenable to QTL identification and breeding in future.

Roots are targets for wheat breeding today and phenotypes such as rapid and long seminal roots and long lateral roots, along with long first leaf growth, are good targets for direct selection in breeding for whole plant establishment. Rapid establishment confers greater ground cover, greater root system soil colonization and improve crop nutrient-use efficiencies (Pang et al., 2013). These root and shoot traits combined are advantageous for contemporary farming systems that do not plough the soil (Watt et al., 2005). Selecting a single trait may not be effective. Targeting two key functional traits (one below and one above ground) with high heritability may serve as more reliable index for trait selection. A platform such as *GrowScreen-PaGe* allows co-selection of root and shoot traits, potentially speeding up introgression of traits and genetic gain within a germplasm enhancement pipeline (Tracy et al., 2020).

### Key Traits for Potential Selection in Breeding

Similar to other studies across plant species, we found the heritability and repeatability were moderate to high (0.30-0.67) for length of first seminal root, lateral root and total root lengths, leaf length, leaf number, and shoot biomass (Table 2, 3). Moderate to high heritability for primary and lateral root length is reported in Arabidopsis (Richard et al., 2011), wheat (Dhanda et al., 2004; Laperche et al., 2006), rice (MacMillan et al., 2006), and soybean (Ao et al., 2010). Rebetzke and Richards (1999) reported that seedling leaf width was highly heritable and had a high genetic correlation with total leaf area in wheat during the vegetative stage. Similarly, Gioia et al. (2016) reported high heritability for leaf number (*H^2^*=0.95) and total root length (*H^2^*=0.93) in rapeseed. Traits with higher heritability are often controlled by fewer genes with larger effects (Tsegaye et al., 2012). Thus, the direct selection of traits with high heritability under strong genetic control enable genetic gain.

### CSIRO MAGIC Parents Grew Faster than NIAB MAGIC Parents

In general, the CSIRO MAGIC parents grew faster, produced larger root systems (mainly through greater total lateral root length), wider seminal root angle (SRA_2cm_) and more plant biomass compared to NIAB MAGIC parents (Figures 3, S2). Differences between the founder parents may be explained by the different origin, genetic background and breeding targets of the populations (Figure 1, Table S1). Four of the CSIRO parents were bred in Australia, possibly contributing genes for traits conferring performance in low rainfed environments, such as rapid and long lateral root growth (Palta and Watt, 2009). NIAB parents come from the UK and France, environments with higher rainfall than Australian wheat growing regions and with higher planting density practices, which may have selected against high total root length (Fradgley et al., 2020; Mackay et al., 2014). Additionally, narrow genetic background during breeding may have resulted in a smaller root system of NIAB parents compared to CSIRO (Camargo et al., 2018; Fradgley et al., 2020; Subbiah et al., 1968). Furthermore, two NIAB parents, Robigus and Soissons, carry the major dwarfing locus, ‘reduced plant height’ (*Rht-B1*), while six other parents carry the plant height reducing locus, *Rht-D1* (Mackay et al., 2014). These dwarfing genes are gibberellic acid insensitive, decrease cell elongation and reduce leaf length and shoot biomass (Botwright et al., 2005; Ellis et al., 2004). In case of CSIRO, two parents Chara7 and Baxter carry *Rht-B1* while Yitpi and Westonia carry the *Rht-D1* (Huang et al., 2012). The effects of dwarfing genes on coleoptile length and plant height of CSIRO parents were reported to be negative (Delhaize et al., 2015; Huang et al., 2012). A number of studies have shown that dwarfing genes varying across MAGIC parents were associated with large reductions in coleoptile length that can impact the seedling establishment (Rebetzke et al. 2007). For example, reduction in coleoptile length was greater for *Rht-D1* than *Rht-B1*. Furthermore, influence of embryo and seed size on early vigour and seedling traits were widely described in wheat (Maydup et al. 2012; Moore and Rebetzke, 2015). Our study did not focus on seed size, though we have taken a great care for seedling uniformity by transplanting similar germinated seed (equal size) of MAGIC parents on germination paper.

The significantly lower lateral root length in slow growing parents compared to fast growing parents (Figure 4A-C, E-G) was a notable finding of this study. The exponential growth of lateral roots in fast growing parents Robigus and AC Barrie led to larger root systems compared to slow growing parents Rialto and G204 Xiaoyan54 (Figure 4D, H). Lateral roots are critical for exploring large volumes of soil for nutrients and water (von Wangenheim et al., 2020). The role of contrasting RSA found in this study can be validated in future with experiments analyzing water and nutrient uptake by slow and fast growing parents, and progeny within the populations.

### Seminal Root Angle was not Correlated to Root System Depth

Although wider seminal root angle (SRA_5cm_) was associated with longer total root length, we did not find a significant correlation with root system depth or width across both sets of MAGIC parents (Figure 6A, B). Rich et al. (2020) recently reported influence of the cultivation systems on formation of root angle in 19 diverse wheat cultivars. They found no significant difference in root angle between first pairs of seminal roots grown in pouches, which was very similar to our findings. However, Rich et al. (2020) found significantly different root angles between second pairs of seminal roots when plants were grown in agar and soil. In contrast, Manschadi et al., (2008); Nakamoto and Oyanagi, (1994); and Oyanagi (1994) reported that a narrow root angle promotes deeper root growth, also in wheat. In those studies, the authors used diverse wheat cultivars which have wider genetic background compared to MAGIC parents analyzed in this study. They were grown in different cultivation systems and measured root angle differently compared to our study. These factors may have resulted the contrasting results. Root angle is affected by the physical, chemical and biological conditions of soil and agar medium (Watt et al., 2006), and this makes it a challenging phenotype for selection in controlled conditions and translation to the field (Rich et al., 2020).

### Biomass Allocation Patterns Vary in Between Fast and Slow Growing Parents

The biomass of roots and shoots were positively correlated (*R*^2^≥0.74) in both sets of MAGIC parents (Figure 7D) suggesting that the root and shoot dry weight are linked and genetically shared the proportion of variance. Our results revealed notable differences in biomass allocation pattern between MAGIC parents expressed as root: shoot ratio (Table 3). Fast growing parents with larger root system (Robigus and AC Barrie) allocated a higher proportion of total biomass into roots, whereas the slow growing parents (Rialto and G204 Xiaoyan54) invested more resources to the shoots at 16 DAT (Table 3). Early allocation of more resources into roots covers the larger volume of ground, facilitates optimal intake of water and nutrient and helps rapid establishment. Our finding further suggests that greater belowground investment also results in larger shoot biomass. For example, the fast growing parents Robigus and AC Barrie revealed both a larger root system and larger leaf length (up to 1.3 times, Supplementary Tables S2 and S3) as compared to small root system parents Rialto and G204 Xiaoyan54. This may indicate a potential higher assimilation of canopy photosynthesis by large parents and supply of carbon to roots after depletion of seed reserves (Palta and Gregory, 1997; Palta et al., 2011). Genotypes with rapid leaf growth were reported to perform better under physical and biological constraints of unploughed soil (Watt et al., 2005). Under resource limited environments, lines with larger root system and leaf area resulted more biomass and yield compared to the lines with low vigour (Zhang et al., 2015).

## CONCLUSION

Root system architecture varies across parents of NIAB and CSIRO MAGIC wheat populations. Faster growing genotypes initially allocated more biomass to roots than shoots, followed by faster lateral root growth and longer leaf length, compared to slower growing genotypes. Lengths of lateral roots and leaves had greatest hertitabilities of all phenotypes measured, and are promising targets for enhancing wheat crop establishment, a critical stage for crop productivity. Phenotyping of progeny of MAGIC parents will be necessary to identify and map QTLs, and dynamic phenotypes can be selected at high throughput using the updated version of *GrowScreen-PaGe* used in this study.

## DATA AVAILABILITY STATEMENT

The datasets generated are available for academic partners for non-commercial purposes upon request sent to the corresponding authors, provided that bilateral terms-of-use agreements can be concluded.

## AUTHOR CONTRIBUTIONS

KAN, CF, and MW designed the study. JL, AG, and SRP implemented the experiments and performed the data analysis. JW, AP, and KH updated the image-processing workflow and AP, SA, and AG designed the new growth container lids and modified the setup. SRP, KAN, and MW worked on interpretation of the data and drafted the manuscript. All authors read and approved the manuscript.

## ACKNOWLEDGMENTS

We thank Carmen Müller, Bernd Kastenholz, and Ann-Katrin Kleinert for their assistance during the experiments, and Sebastian Steinemann for his help in setting up the experimental design. Furthermore, we thank Natalie Wuyts and Jose Correa for support in data (*R* - program) analysis. Michelle Watt holds the Adrienne Clarke Professorial Chair of Botany, which is supported through the University of Melbourne Botany Foundation. We are grateful for the provision of MAGIC parents by NIAB and CSIRO.

## SUPPLEMENTARY MATERIALS

The supplementary materials are included with this article.

**Supplementary Figure S1.** Workflow of the phenotyping system *GrowScreen-PaGe*. The wheat plants are grown in grey opaque containers.

**Supplementary Figure S2.** Variation for root angle measured between the two outmost seminal roots at a distance of 2 cm (SRA_2cm_) at A) 2 days after transplanting (DAT), B) 9 DAT and C) 16 DAT

**Supplementary Table S1.** List of MAGIC wheat parents used in this study, their origin, pedigree, and trait attributes.

**Supplementary Table S2.** Phenotyping variation for root and shoot traits of NIAB parents over time, mean values are shown.

**Supplementary Table S3.** Phenotyping variation for root and shoot traits of CSIRO parents over time, mean values are shown.

**Supplementary Table S4.** Correlations (*R^2^*) between root and shoot traits of MAGIC wheat parents. Root and shoot dry weight were measured at 16 days after transplanting.

**Supplementary Table S5.** Coefficient of variance (CV) across NIAB MAGIC wheat parents over time.

**Supplementary Table S6.** Coefficient of variance (CV) across CSIRO MAGIC wheat parents over time.

