## Supplementary materials for "Variation in root system architecture among the founder parents of two 8-way MAGIC wheat populations for selection in breeding"

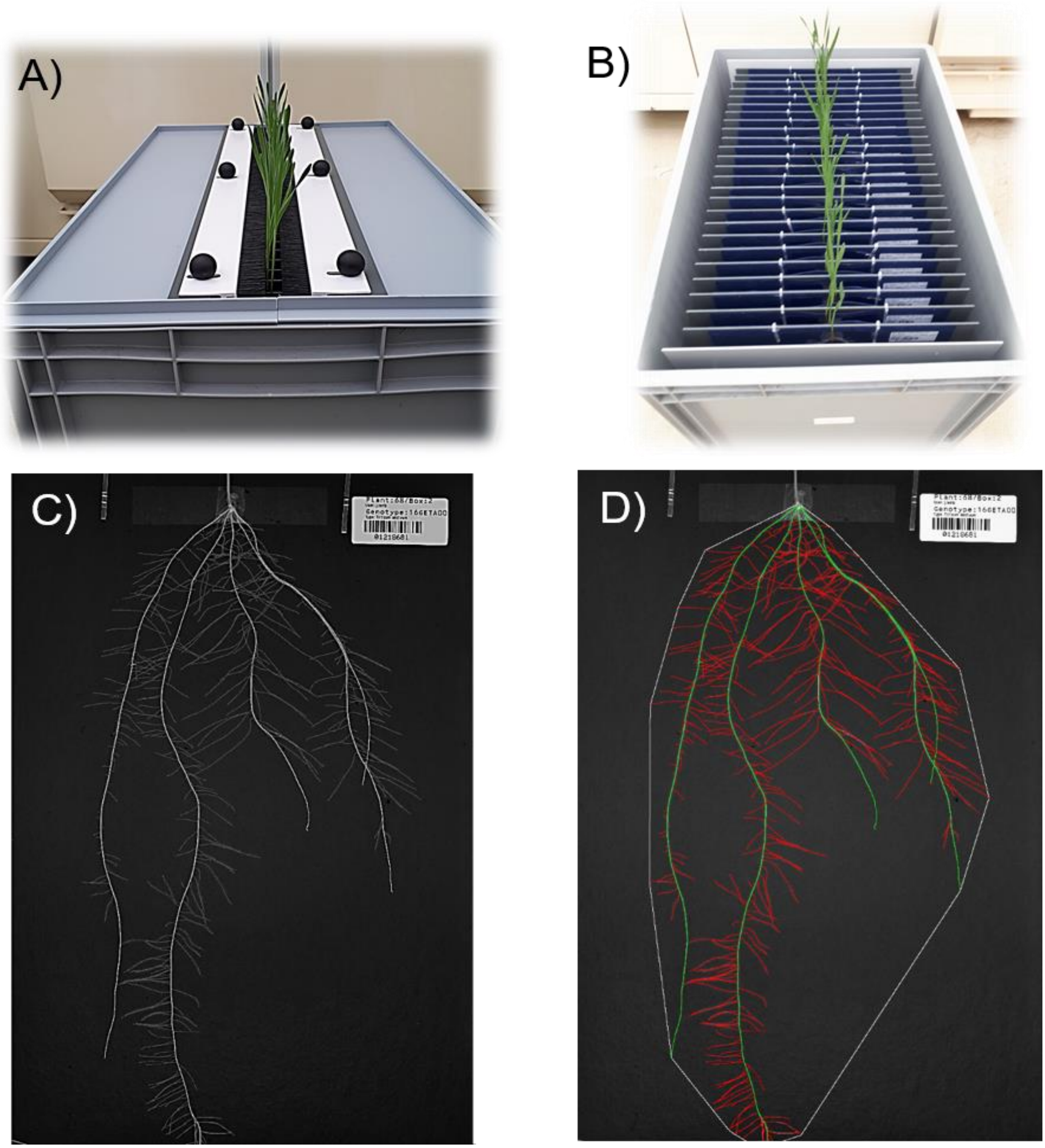


**Supplementary Figure S1**. Workflow of the phenotyping system "*GrowScreen-PaGe*". The wheat plants are grown in grey opaque containers. To prevent light reaching the roots the containers are covered by a custom-made lid with an adjustable black strip brushes in the middle (A). In each box 34 plants are grown (B, for better illustration the lid has been removed). A typical original root image captured (C) and analyzed by an image-processing workflow (D; seminal roots are shown in green and lateral roots in red). The barcode on each images enables the automatic identification of the PlantID. The images (C, D) show the root system of genotype Xi-19 16 days after transplanting.

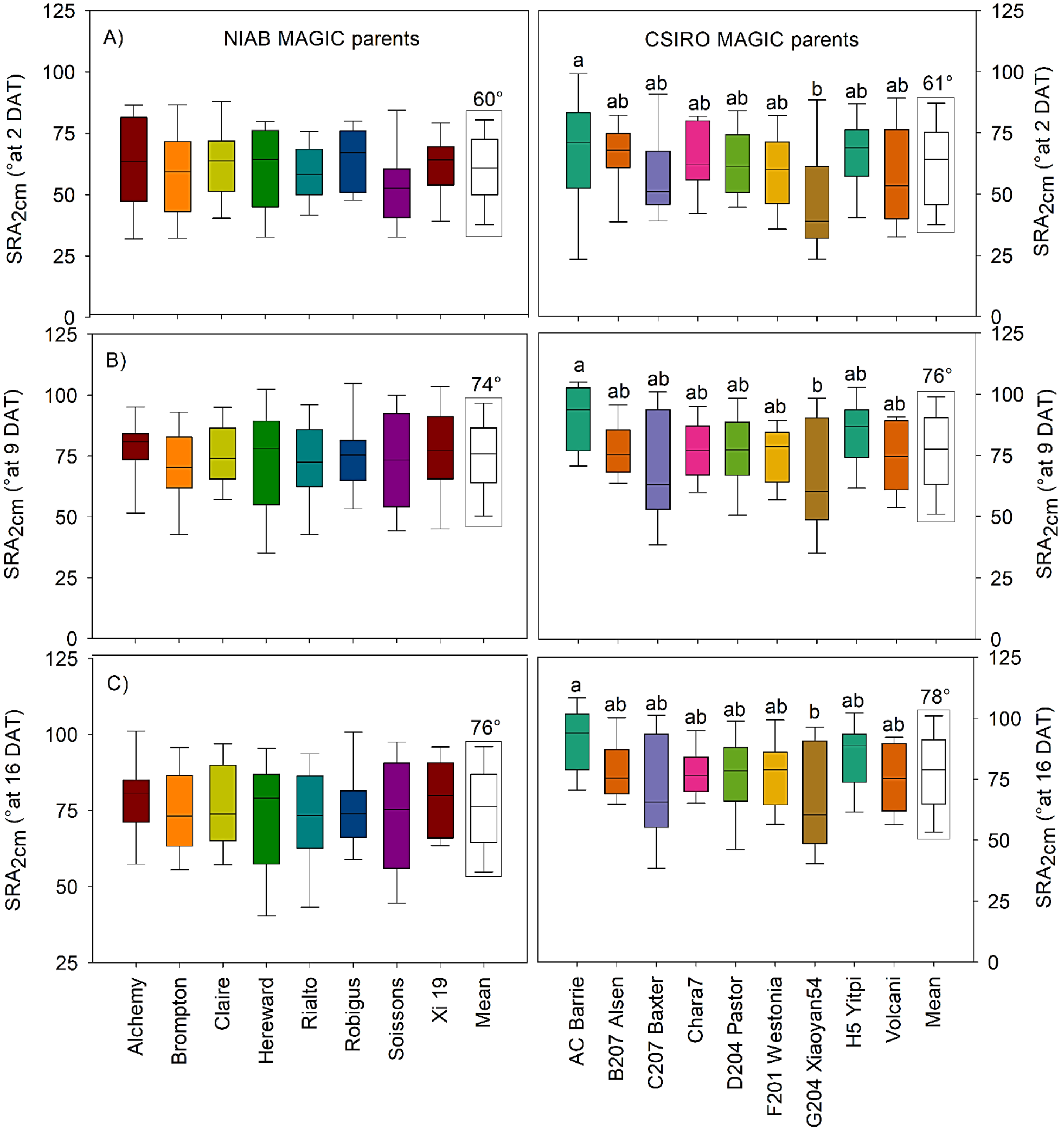


**Supplementary Figure S2**. Variation for root angle measured between the two outmost seminal roots at a distance of 2 cm ($SRA_{2cm}$) at A) 2 days after transplanting (DAT), B) 9 DAT and C) 16 DAT. The boxes indicate the 25$^{th}$ and 75$^{th}$ percentile of the distribution, the 'whiskers' the 10$^{th}$ and 90$^{th}$ percentile and the lines in the middle of the box the median value. No significant differences were found at $P < 0.05$, n = 24 according to one way ANOVA analysis. Boxes highlighted with rectangular black line indicates mean seminal root angle (SRA) across NIAB and CSIRO wheat parents.

Supplementary Table S1. List of MAGIC wheat parents used in this study, their origin, pedigree, and trait attributes

| CName | MAGIC founder lines | Pedigree | Trait attributes | Source/Breeder | Ploidy/planting | Generations | Traits/qtls | References |
|---|---|---|---|---|---|---|---|---|
| Alchemy | NIAB-eight parents | Claire x (Consort x Woodstock) | High yield, disease resistance, breeding use, soft wheat | Limagrain UK Ltd, UK | Hexaploid/winter | 1091/F7 (90K SNP) | Yield, flowering time, plant height, yellow rust, fusarium diseases, mildew, awn presence | Machekey et al. (2014) and Scutari et al. (2014) |
| Brompton | | CWW 92.1 x Caxton | Hard feed, 1BL/1RS, resistant to OWBM | Elsoms Seeds Ltd, UK | | | | |
| Claire | | Wasp x Flame | Soft biscuit/distilling, slow apical development | Limagrain UK Ltd, UK | | | | |
| Hereward | | Norman sib x Disponent | High-quality benchmark-1 bread-making | RAGT seeds Ltd, UK | | | | |
| Rialto | | Haven x Fresco | Moderate bread-making, 1BL/1RS | RAGT seeds Ltd, UK | | | | |
| Robigus | | Z836 x 1366 (PUTCH) | Exotic introgression, disease resistance, breeding use, Rht-B1 | KWS UK Ltd, UK | | | | |
| Soissons | | Iena (Jena) x HN35 | Bread-making quality, early flowering, Rht-B1 | Maison Florimond Desprez, France | | | | |
| Xi 19 | | (Cadenza x Rialto) x Cadenza | Bread-making quality, facultative type, breeding use | Limagrain UK Ltd, UK | | | | |
| AC Barrie | CSIRO-eight parents | Neepawa/ Columbus//BW90 | Hard red spring wheat, high yield and high protein content, and resistant to lodging | Canada | Hexaploid/spring | 2099/F6 (90K SNP+GBS) | Coleoptile weight, size and length, root length, root hairs and rhizosheath, and plant height | Hunag et al. (2012), Cavanagh et al. (2013), Rebetzke et a.(2014), and http://www.bioplatforms.com/variety-sequencing/ |
| B207 Alsen | | Grandin/Glupro/5/Sumai3/Wheaton//Grandin/4/Grandin/3/IAS20*4/H567.71//Amidon | Hard red spring wheat, moderate to stem and leaf rust and Fusarium head blight disease | USA | | | | |
| C207 Baxter | CSIRO- four/eight parents- | QT2327/Cook//QT2804 | Resistance to RLN, semi draft | (GDRC), Australia | | | | |
| Chara7 | CSIRO-four parents | COOK*2/MILLEWA//TM56/PAVON/CONDOR | High yielding, broadly adapted wheat, moderately tolerance to acidic soils and CCN | (GDRC), Australia | | | | |
| D204 Pastor | CSIRO-eight parents | PFAU/SERI-82(CID-93300//BOBWHITE | Semi-dwarf in height, moderate resistance to rust | CIMMYT, Mexico | | | | |
| F201 Westonia | CSIRO-four/eight parents | SPICA/TIMGALEN//TOSCA/3/CRANBROOK//BOB-WHITE*2/JACUP | Good aluminium tolerance, susceptible to stem rust (SVS) and stripe rust | (GDRC), Australia | | | | |
| G204 Xiaoyan54 | CSIRO-eight parents | (M)ST-2422-464/XIAOYAN-86 | High quality noodle wheat, resistant to stripe rust | China | | | | |
| H5 Yitpi | CSIRO- four/eight parents | C8MMC8HMM/Frame | Large grain, long coleoptile, tolerant to CCN and boron. susceptible to stem rust (S) and yellow spot (SVS) | (GDRC), Australia | | | | |
| Volcani | CSIRO-eight parents | V761-28-J4-B2-NZ8 | Good milling, strong dough and good baking properties, contains the grain protein content gene and Yr36 resistance to stem, stripe and leaf rust | Isreal | | | | |

* CName common name, MAGIC multiparent advance generations of inter-cross, NIAB The National Institute of Agricultural Botany; CSIRO Commonwealth Scientific and Industrial Research Organisation; qtl quantittative trait orange wheat blossom midge, loci, OWBM Rht reduced height, RLN root lession nematodes, CCN cereal cyst nematodes

Supplementary Table S2. Phenotyping variation for root and shoot traits of NIAB parents over time, mean values are shown

| Traits | Time points | NIAB-wheat-MAGIC-parent lines | | | | | | | | Mean |
|---|---|---|---|---|---|---|---|---|---|---|
| | | **Alchemy** | **Brompton** | **Claire** | **Hereward** | **Rialto** | **Robigus** | **Soissons** | **Xi 19** | |
| First seminal root (mm) | 1 DAG | 1.63ab | 1.5ab | 1.7ab | 1.66ab | 1a | 2b | 1.37ab | 1.66ab | 1.565 |
| Seminal root length (cm) | 2 DAT | 14.61ab | 14.73ab | 14.23ab | 13.67ab | 12.05a | 17.51b | 11.76a | 12.71a | 13.909 |
| | 9 DAT | 83.27b | 82.7b | 76.59ab | 71.07ab | 65.98a | 85.27b | 76.1ab | 68.32a | 76.163 |
| | 16 DAT | 130.75b | 119.19b | 118.83ab | 111.17ab | 109.03a | 122.52b | 132.43b | 102.98ab | 119.613 |
| Lateral root length (cm) | 2 DAT | NA | | | | | | | | |
| | 9 DAT | 28.84ab | 36.06b | 37.51bc | 18.95a | 18.81a | 49.20c | 28.23ab | 24.02ab | 30.20 |
| | 16 DAT | 284.19ab | 365.92b | 346.56ab | 209.29a | 178.80a | 364.89b | 279.08ab | 299.94ab | 291.08 |
| Total root length (cm) | 2 DAT | 14.61ab | 14.73ab | 14.23ab | 13.67ab | 12.05a | 17.51b | 11.76a | 12.71a | 13.91 |
| | 9 DAT | 112.12b | 118.76bc | 114.10b | 90.031a | 84.79a | 134.48c | 104.33ab | 92.34a | 106.38 |
| | 16 DAT | 414.95b | 474.12b | 465.40b | 320.46a | 287.84a | 487.41b | 411.52b | 402.93ab | 410.71 |
| Root system depth (cm) | 2 DAT | 4.99 | 4.87 | 5.20 | 5.07 | 4.57 | 6.53 | 4.32 | 4.56 | 5.02 |
| | 9 DAT | 21.00ab | 20.89ab | 21.83ab | 19.69ab | 19.00ab | 24.43b | 20.58ab | 18.34a | 20.73 |
| | 16 DAT | 32.42 | 31.97 | 33.31 | 31.06 | 31.75 | 32.93 | 30.88 | 29.76 | 31.76 |
| Root system width (cm) | 2 DAT | 3.65 | 3.76 | 4.31 | 3.78 | 3.34 | 4.73 | 2.89 | 3.50 | 3.75 |
| | 9 DAT | 14.09ab | 14.90ab | 18.02b | 14.72ab | 13.38ab | 14.58ab | 11.59a | 13.93ab | 14.41 |
| | 16 DAT | 19.03 | 19.79 | 22.06 | 18.99 | 18.19 | 18.26 | 17.30 | 18.47 | 19.01 |
| Root angle at 2 cm distance (⁰) | 2 DAT | 62.79 | 58.09 | 62.53 | 60.06 | 58.69 | 64.67 | 53.86 | 61.05 | 60.00 |
| | 9 DAT | 78.12 | 70.57 | 74.94 | 73.54 | 71.75 | 75.37 | 73.30 | 77.06 | 74.00 |
| | 16 DAT | 79.29 | 74.95 | 75.98 | 74.09 | 72.68 | 75.32 | 73.49 | 78.30 | 76.00 |
| Root angle at 5 cm distance (⁰) | 2 DAT | NA | | | | | | | | |
| | 9 DAT | 62.38 | 62.59 | 69.18 | 61.33 | 54.64 | 61.44 | 54.19 | 60.80 | 61.00 |
| | 16 DAT | 63.68 | 63.53 | 70.03 | 63.43 | 57.28 | 64.18 | 61.30 | 64.97 | 64.00 |
| Convex hull area (cm²) | 2 DAT | 11.43ab | 9.74a | 12.63ab | 11.08ab | 8.97a | 18.33b | 8.042a | 9.47a | 11.21 |
| | 9 DAT | 192.90b | 178.04ab | 227.33b | 158.37ab | 157.35ab | 208.49b | 149.46a | 148.02a | 177.50 |
| | 16 DAT | 416.80b | 414.85b | 475.88c | 368.25a | 387.96ab | 406.86b | 378.15ab | 354.26a | 400.38 |
| Leaf length (cm) | 16 DAT | 14.00ab | 14.50b | 13.02ab | 11.10a | 12.23a | 14.96b | 17.38c | 12.66a | 13.74 |
| Number of leaves | 16 DAT | 2.07 | 2.44 | 2.13 | 1.94 | 1.73 | 2.50 | 2.31 | 2.13 | 2.15 |
| Root dry weight (g) | 16 DAT | 0.022b | 0.027b | 0.021ab | 0.017a | 0.018a | 0.023b | 0.024b | 0.020ab | 0.0218 |
| Shoot dry weight (g) | 16 DAT | 0.0209ab | 0.0244b | 0.0197ab | 0.0175a | 0.0186a | 0.0216ab | 0.023b | 0.0178a | 0.0205 |

* MAGIC Multiparent advanced generation intercross; NIAB The National Institute of Agricultural Botany; mean data presented; different letters are signifcantly different at *p=0.05* , n=24; a < b < c; NA data not available; DAG days after germination and DAT days after transplanting

Supplementary Table S3. Phenotyping variation for root and shoot traits of CSIRO parents over time, mean values are shown.

| **Traits** | **Time points** | **CSIRO-wheat-MAGIC-parent lines** | | | | | | | | | **Mean** |
|---|---|---|---|---|---|---|---|---|---|---|---|
| | | **AC Barrie** | **B207 Alsen** | **C207** | **Chara7** | **D204** | **F201 Westonia** | **G204 Xiaoyan54** | **H5 Yitpi** | **Volcani** | |
| First seminal root (mm) | 1 DAG | 1.37a | 1.75a | 1.20a | 2.41ab | 2.16ab | 3.62b | 1.37a | 1.95a | 1.33a | 1.91 |
| Seminal root length (cm) | 2 DAT | 19.32 | 20.91 | 15.77 | 15.70 | 17.60 | 18.79 | 14.29 | 17.61 | 15.29 | 17.25 |
| | 9 DAT | 94.63b | 106.12b | 85.12ab | 81.21a | 91.65b | 86.89ab | 68.47a | 88.17ab | 85.76ab | 87.56 |
| | 16 DAT | 137.65ab | 149.92b | 133.72ab | 124.65ab | 136.22b | 126.23ab | 107.12a | 137.1b | 140.12b | 132.53 |
| Lateral root length (cm) | 2 DAT | NA | | | | | | | | | |
| | 9 DAT | 56.13b | 31.66ab | 36.39ab | 45.29b | 43.51ab | 35.91ab | 44.11ab | 49.03b | 21.66a | 40.42 |
| | 16 DAT | 554.15c | 416.54b | 330.49ab | 399.95b | 392.93b | 363.27b | 249.88a | 390.63b | 312.72b | 378.96 |
| Total root length (cm) | 2 DAT | 19.32 | 20.91 | 15.77 | 15.70 | 17.60 | 18.79 | 14.29 | 17.61 | 15.29 | 17.25 |
| | 9 DAT | 150.77b | 137.78b | 121.51a | 126.51ab | 135.17ab | 122.81a | 112.59a | 137.20b | 107.43 | 127.98 |
| | 16 DAT | 691.81b | 566.47b | 464.21ab | 524.61b | 529.16b | 489.50ab | 357.00a | 527.74b | 452.85ab | 511.49 |
| Root system depth (cm) | 2 DAT | 5.68 | 5.90 | 5.19 | 5.38 | 5.78 | 6.33 | 5.33 | 6.85 | 5.17 | 5.74 |
| | 9 DAT | 23.31ab | 24.40ab | 20.99ab | 22.94ab | 22.33ab | 24.03b | 21.60ab | 21.90ab | 20.92a | 22.50 |
| | 16 DAT | 33.77 | 32.93 | 32.78 | 32.71 | 32.43 | 32.57 | 32.71 | 32.50 | 33.00 | 32.82 |
| Root system width (cm) | 2 DAT | 4.58 | 4.26 | 3.51 | 4.41 | 4.71 | 3.91 | 3.14 | 4.20 | 3.63 | 4.04 |
| | 9 DAT | 17.10b | 16.43b | 13.44ab | 14.28ab | 14.70ab | 15.13ab | 11.10a | 14.88ab | 12.34a | 14.38 |
| | 16 DAT | 21.52 | 20.15 | 19.06 | 18.91 | 19.84 | 20.67 | 15.83 | 19.75 | 18.87 | 19.40 |
| Root angle at 2 cm distance (⁰) | 2 DAT | 69a | 66ab | 58ab | 64ab | 63ab | 59ab | 47b | 67a | 58ab | 61.00 |
| | 9 DAT | 89a | 76ab | 72ab | 77ab | 77ab | 75ab | 64b | 84ab | 73ab | 76.00 |
| | 16 DAT | 90a | 79ab | 73ab | 78ab | 76ab | 76ab | 63b | 84ab | 75ab | 78.00 |
| Root angle at 5 cm distance (⁰) | 2 DAT | NA | | | | | | | | | |
| | 9 DAT | 68a | 59ab | 59ab | 55ab | 65ab | 60ab | 46b | 61ab | 53ab | 58.00 |
| | 16 DAT | 68 | 58 | 59 | 57 | 64 | 58 | 48 | 62 | 56 | 59.00 |
| Convex hull area (cm²) | 2 DAT | 17.08b | 15.57ab | 12.19ab | 13.57ab | 16.84ab | 14.20ab | 9.33a | 17.56b | 11.59a | 14.22 |
| | 9 DAT | 242.33 | 228.58 | 178.23 | 190.69 | 183.96 | 198.62 | 136.82 | 198.27 | 164.52 | 191.34 |
| | 16 DAT | 502.98 | 448.82 | 413.66 | 404.42 | 415.86 | 427.55 | 334.49 | 456.67 | 405.58 | 423.34 |
| Leaf length (cm) | 16 DAT | 21.49b | 20.32b | 14.95a | 15.70ab | 16.69ab | 16.65ab | 10.26a | 17.26ab | 15.96ab | 16.59 |
| Number of leaves | 16 DAT | 2.54 | 3.04 | 3.00 | 3.00 | 2.88 | 2.83 | 2.74 | 2.79 | 2.38 | 2.80 |
| Root dry weight (g) | 16 DAT | 0.0329b | 0.0302b | 0.0241ab | 0.0260ab | 0.026ab | 0.026ab | 0.0178a | 0.0257ab | 0.0277b | 0.026 |
| Shoot dry weight (g) | 16 DAT | 0.0293b | 0.0311b | 0.0214a | 0.0228ab | 0.0234ab | 0.0236ab | 0.0180a | 0.0256ab | 0.0243ab | 0.024 |

* MAGIC Multiparent advanced generation intercross; CSIRO Commonwealth Scientific and Industrial Research Organisation; mean data presented; different letters are signifcantly different at $p=0.05$ , n=24; a < b < c; NA data not available; DAG days after germination and DAT days after transplanting.

Supplementary Table S4. Correlations (R2) between root and shoot traits of MAGIC wheat parents. Root and shoot dry weight were measured at 16 days after transplanting.

| **Cname** | **MAGIC founder parents** | **Correlation between** | | | |
|---|---|---|---|---|---|
| | | **SRL and LRL** | **RDW and TRL** | **SDW and TRL** | **RDW and SDW** |
| Alchemy | NIAB | 0.245 | 0.402 | 0.260 | 0.761 |
| Bromptom | | 0.135 | 0.689 | 0.747 | 0.809 |
| Claire | | 0.159 | 0.309 | 0.249 | 0.654 |
| Hereward | | 0.219 | 0.619 | 0.609 | 0.785 |
| Rialto | | 0.339 | 0.499 | 0.586 | 0.762 |
| Robigus | | 0.112 | 0.602 | 0.627 | 0.627 |
| Soissions | | 0.376 | 0.393 | 0.872 | 0.872 |
| Xi-19 | | 0.325 | 0.591 | 0.771 | 0.772 |
| AC Barrie | CSIRO | 0.003 | 0.693 | 0.647 | 0.487 |
| B207 Alsen | | 0.269 | 0.599 | 0.518 | 0.852 |
| C207 Baxter | | 0.055 | 0.668 | 0.449 | 0.655 |
| Chara7 | | 0.004 | 0.784 | 0.627 | 0.685 |
| D204 Pastor | | 0.008 | 0.663 | 0.439 | 0.426 |
| F201 Westonia | | 0.351 | 0.770 | 0.887 | 0.818 |
| G204 Xiaoyan54 | | 0.061 | 0.731 | 0.699 | 0.898 |
| H5 Yitpi | | 0.378 | 0.616 | 0.740 | 0.865 |
| Volcani | | 0.037 | 0.585 | 0.571 | 0.686 |

*Cname common name, MAGIC multiparent advance generations of inter-cross, NIAB The National Institute of Agricultural Botany; CSIRO Commonwealth Scientific and Industrial Research Organisation; SRL seminal root length, TRL total root length, RDW root dry weight, SDW shoot dry weight, root and shoot dry weight were measured at 16 days after transplanting

Supplementary Table S5. Coefficient of variance (CV) across NIAB MAGIC wheat parents over time.

| Traits | Time points | NIAB-Wheat MAGIC parents | | | | | | | |
|---|---|---|---|---|---|---|---|---|---|
| | | **Alchemy** | **Brompton** | **Claire** | **Hereward** | **Rialto** | **Robigus** | **Soissons** | **Xi 19** |
| First seminal root (mm) | 1 DAG | 44 | 47 | 47 | 49 | 20 | 43 | 45 | 39 |
| | 2 DAT | 37 | 26 | 32 | 23 | 34 | 19 | 29 | 30 |
| Seminal root length (cm) | 9 DAT | 32 | 24 | 19 | 24 | 31 | 19 | 30 | 29 |
| | 16 DAT | 26 | 22 | 16 | 28 | 19 | 20 | 27 | 23 |
| | 2 DAT | NA | | | | | | | |
| Lateral root length (cm) | 9 DAT | 49 | 42 | 43 | 51 | 42 | 40 | 62 | 40 |
| | 16 DAT | 39 | 32 | 34 | 40 | 48 | 30 | 44 | 35 |
| | 2 DAT | 34 | 27 | 24 | 23 | 28 | 21 | 35 | 32 |
| Total root length (cm) | 9 DAT | 29 | 26 | 23 | 27 | 33 | 26 | 37 | 34 |
| | 16 DAT | 38 | 27 | 27 | 35 | 46 | 23 | 43 | 30 |
| | 2 DAT | 33 | 28 | 20 | 26 | 25 | 16 | 38 | 28 |
| Root system depth (cm) | 9 DAT | 28 | 24 | 12 | 15 | 20 | 13 | 31 | 21 |
| | 16 DAT | 15 | 16 | 4 | 13 | 11 | 7 | 16 | 18 |
| | 2 DAT | 38 | 34 | 31 | 34 | 42 | 29 | 44 | 41 |
| Root system width (cm) | 9 DAT | 33 | 26 | 24 | 28 | 35 | 25 | 41 | 38 |
| | 16 DAT | 21 | 20 | 12 | 22 | 26 | 20 | 32 | 27 |
| | 2 DAT | NA | | | | | | | |
| Seminal root angle (2cm, º) | 9 DAT | 32 | 29 | 29 | 31 | 47 | 41 | 31 | 38 |
| | 16 DAT | 28 | 33 | 29 | 33 | 40 | 31 | 37 | 45 |
| | 2 DAT | 43 | 38 | 42 | 47 | 47 | 41 | 57 | 43 |
| Convex hull area (cm²) | 9 DAT | 44 | 36 | 30 | 40 | 46 | 32 | 42 | 34 |
| | 16 DAT | 30 | 28 | 20 | 29 | 36 | 30 | 30 | 25 |
| Leaf length (cm) | 16 DAT | 26 | 15 | 19 | 24 | 18 | 19 | 20 | 20 |
| Number of leaves | 16 DAT | 25 | 18 | 38 | 29 | 26 | 25 | 33 | 40 |
| Root dry weight (g) | 16 DAT | 39 | 29 | 20 | 34 | 41 | 23 | 41 | 38 |
| Shoot dry weight (g) | 16 DAT | 38 | 30 | 26 | 36 | 40 | 25 | 34 | 39 |
| Total biomass (g) | 16 DAT | 37 | 28 | 22 | 35 | 40 | 21 | 38 | 38 |

* NIAB The National Institute of Agricultural Botany; MAGIC Multiparent advanced generation intercross; CV coefficient of variance ; n=24; NA data not available; DAG days after germination, DAT Days after transplanting

Supplementary Table S6. Coefficient of variance (CV) across CSIRO MAGIC wheat parents over time.

| **Traits** | **Time points** | **CSIRO-MAGIC wheat parents** | | | | | | | | |
|---|---|---|---|---|---|---|---|---|---|---|
| | | **AC Barrie** | **B207 Alsen** | **C207 Baxter** | **Chara7** | **D204 Pastor** | **F201 Westonia** | **G204 Xiaoyan54** | **H5 Yitpi** | **Volcani** |
| First seminal root (mm) | 1 DAG | 41 | 36 | 35 | 34 | 45 | 46 | 47 | 48 | 31 |
| Seminal root length (cm) | 2 DAT | 29 | 18 | 33 | 32 | 25 | 26 | 24 | 45 | 32 |
| | 9 DAT | 22 | 18 | 23 | 16 | 18 | 24 | 19 | 33 | 22 |
| | 16 DAT | 19 | 18 | 16 | 20 | 17 | 25 | 18 | 21 | 18 |
| Lateral root length (cm) | 2 DAT | NA | | | | | | | | |
| | 9 DAT | 62 | 54 | 63 | 54 | 44 | 49 | 58 | 52 | 49 |
| | 16 DAT | 31 | 36 | 51 | 43 | 38 | 40 | 48 | 43 | 52 |
| Total root length (cm) | 2 DAT | 29 | 15 | 33 | 32 | 25 | 23 | 24 | 25 | 27 |
| | 9 DAT | 23 | 23 | 33 | 27 | 23 | 33 | 29 | 35 | 28 |
| | 16 DAT | 24 | 29 | 35 | 33 | 27 | 39 | 34 | 36 | 31 |
| Root system depth (cm) | 2 DAT | 25 | 14 | 24 | 27 | 25 | 19 | 20 | 45 | 24 |
| | 9 DAT | 16 | 12 | 15 | 24 | 12 | 19 | 16 | 18 | 17 |
| | 16 DAT | 3 | 8 | 7 | 7 | 8 | 10 | 9 | 9 | 7 |
| Root system width (cm) | 2 DAT | 36 | 28 | 43 | 40 | 36 | 43 | 49 | 32 | 32 |
| | 9 DAT | 24 | 28 | 26 | 32 | 17 | 24 | 30 | 20 | 30 |
| | 16 DAT | 14 | 19 | 27 | 22 | 12 | 17 | 26 | 20 | 22 |
| Seminal root angle (2cm, º) | 2 DAT | NA | | | | | | | | |
| | 9 DAT | 37 | 30 | 43 | 45 | 34 | 42 | 45 | 37 | 42 |
| | 16 DAT | 39 | 38 | 49 | 45 | 34 | 37 | 42 | 33 | 34 |
| Convex hull area (cm²) | 2 DAT | 41 | 33 | 45 | 35 | 43 | 48 | 47 | 50 | 47 |
| | 9 DAT | 32 | 41 | 45 | 43 | 36 | 30 | 43 | 28 | 43 |
| | 16 DAT | 22 | 29 | 28 | 27 | 23 | 19 | 28 | 26 | 28 |
| Leaf length (cm) | 16 DAT | 9 | 13 | 17 | 8 | 10 | 12 | 18 | 19 | 18 |
| Number of leaves | 16 DAT | 10 | 4 | 0 | 0 | 9 | 10 | 15 | 11 | 15 |
| Root dry weight (g) | 16 DAT | 19 | 22 | 27 | 23 | 18 | 33 | 32 | 34 | 27 |
| Shoot dry weight (g) | 16 DAT | 19 | 29 | 26 | 22 | 24 | 32 | 33 | 28 | 26 |
| Total biomass (g) | 16 DAT | 13 | 16 | 25 | 20 | 20 | 32 | 32 | 22 | 26 |

* CSIRO Commonwealth Scientific and Industrial Research Organisation; MAGIC Multiparent advanced generation intercross; CV coefficient of variance among replications, n=24; and NA data not available; DAG days after germination, DAT Days after transplanting.
